# Morphine selectively promotes glutamate release from glutamatergic terminals of projection neurons from medial prefrontal cortex to dopamine neurons of ventral tegmental area

**DOI:** 10.1101/133090

**Authors:** Li Yang, Ming Chen, Ping Zheng

**Author notes:** Corresponding author: Ping Zheng, PhD, State Key Laboratory of Medical Neurobiology, Shanghai Medical College, Fudan University, 138 Yixueyuan Road, Shanghai 200032, People’s Republic of China.

## Abstract

Recently, we found that morphine promoted presynaptic glutamate release of dopamine (DA) neurons in the ventral tegmental area (VTA), which constituted the main mechanism for morphine-induced increase in VTA-DA neuron firing and related behaviors (Chen et al., 2015). However, what source of presynaptic glutamate release of DA neurons in the VTA is promoted by morphine remains unknown. To address this question, we used optogenetic strategy to selectively activate glutamatergic inputs from different projection neurons and then observed the effect of morphine on them. The result shows that morphine promotes glutamate release from glutamatergic terminals of projection neurons from the medial prefrontal cortex (mPFC) to VTA DA neurons, but has no effect on that from the basolateral amygdala (BLA) or the lateral hypothalamus (LH) to VTA DA neurons, and the inhibition of glutamatergic projection neurons from the mPFC to the VTA significantly reduces morphine-induced increase in locomotor activity of mice.

## Introduction

Morphine-induced addictive behaviors are strongly dependent on the activation of dopamine (DA) neurons of the ventral tegmental area (VTA) (Wise, 1989, Gardner, 2011, Luscher and Malenka, 2011). One previously reported mechanism for morphine to activate VTA-DA neurons is the disinhibition model of VTA-DA neurons (Johnson and North, 1992, Kalivas, 1993, White, 1996). Recently, we found that morphine could promote presynaptic glutamate release of VTA-DA neurons, which constituted the main mechanism for morphine-induced increase in VTA-DA neurons firing and related behaviors (Chen et al., 2015). However, what source of presynaptic glutamate release of DA neurons in the VTA is promoted by morphine remains unknown.

It has been known that DA neurons of the VTA receive glutamatergic inputs from the medial prefrontal cortex (mPFC), the bilateral amygdala (BLA) and the lateral hypothalamus (LH) (Stuber et al., 2012, Li et al., 2013). To study which glutamatergic inputs onto DA neurons of the VTA are modulated by morphine, we used optogenetic strategy to selectively activate glutamatergic inputs from these different projection neurons and then observed the effect of morphine on them. We also studied the contribution of morphine-induced increase in presynaptic glutamate release of DA neurons in the VTA from specific brain region to morphine-induced increase in locomotor activity of mice, which is an index of enhanced DA function in the VTA (Jalabert et al., 2011).

## Results

### Morphine promotes glutamate release from glutamatergic terminals of projection neurons from the mPFC to VTA DA neurons, but has no effect on that from the BLA or the LH to VTA DA neurons

To study whether morphine promoted glutamate release from glutamatergic terminals of projection neurons from the mPFC, the BLA or the LH to VTA DA neurons, we used a pair of optical pulses at intervals of 50 ms at 0.1 Hz to evoke excitatory postsynaptic currents (EPSC) in VTA DA neurons from these different brain areas and then observed the effect of morphine on paired-pulse ratio (PPR) of blue light (470 nm)-evoked EPSCs pulses (intervals of 50 ms at 0.1 Hz). PPR, measured as the ratio of EPSC amplitude in response to two successive stimulation pulses, reflects presynaptic release probability; a lower PPR correlates with higher release probability (Zucker and Regehr, 2002, Li et al., 2013). Left panel of Fig. 1a showed the expression of ChR2–mCherry in the mPFC at 6 weeks after the injection of the virus into the mPFC and right panel of Fig. 1a showed the expression of ChR2-mCherry in the VTA at 6 weeks after the injection of the virus into the mPFC. Left panel of Fig. 1b showed typical original traces of the first and second EPSCs before and after morphine (10 μM) as well as superimposed traces of the first and second EPSCs normalized to the amplitude of the first EPSC in DA neurons of the VTA. The average first EPSC was increased by 50.33 ± 10.39 % (n = 7, from 5 animals), whereas the average second EPSC was increased by 8.03 ± 13.71 % (n = 7, from 5 animals) at 15 min after morphine. So the superimposition of the two traces normalized to the first EPSC showed that PPR significantly decreased after morphine (left panel of Fig. 1b). The average PPR decreased from 1.01 ± 0.07 before morphine to 0.72 ± 0.1 at 15 min after morphine (right panel of Fig. 1b, n = 7, from 5 animals, paired t test, P = 0.0278, compared to control before morphine). However, morphine (10 μM) had no significant effect on PPR evoked by optical stimulation of ChR2-mCherry-positive excitatory terminals from the BLA or the LH to VTA DA neurons. The average PPR was 0.77 ± 0.1 before morphine and 0.8 ± 0.06 at 15 min after morphine in BLA-to-VTA group (right panel of Fig. 2b, n = 11, from 8 animals, paired t test, P = 0.8239, compared to control before morphine) and the average PPR was 0.84 ± 0.07 before morphine and 0.87 ± 0.08 at 15 min after morphine in LH-to-VTA group (right panel of Fig. 3b, n = 7, from 5 animals, paired t test, P = 0.8082, compared to control before morphine). These results suggest that morphine can promote glutamate release from glutamatergic terminals of projection neurons from the mPFC to VTA DA neurons, but has no effect on that from the BLA or the LH to VTA DA neurons.

**Figure 1.**
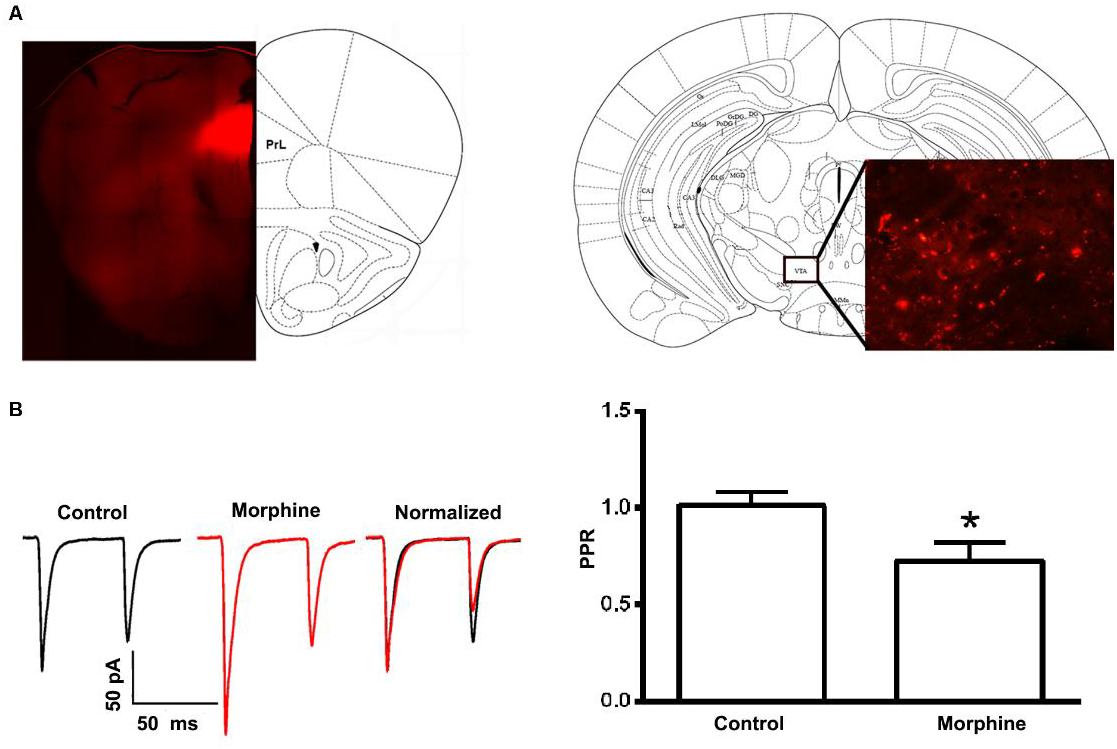
Effect of morphine on paired-pulse ratio (PPR) of light-evoked EPSC in DA neurons of the VTA from the mPFC. (a) Left panel: image of coronal brain slice showing expression of ChR2-mCherry (red color) in the mPFC at 6 weeks after the injection of the virus into the mPFC. Right panel: image of coronal brain slice showing expression of ChR2-mCherry in the VTA at 6 weeks after the injection of the virus into the mPFC. (b) Left panel: typical original traces of the first and second EPSCs evoked by a pair of blue light pulses (470 nm, intervals of 50 ms at 0.1 Hz) before and after morphine (10 μM) as well as superimposed traces of the first and second EPSCs normalized to the amplitude of the first EPSC in DA neurons of the VTA. Right panel: average PPR evoked by a pair of blue light pulses (470 nm, intervals of 50 ms at 0.1 Hz) before and after morphine (10 μM) in VTA DA neurons from the mPFC (n = 7 cells from five mice, P < 0.05, compared to control before morphine).

**Figure 2.**
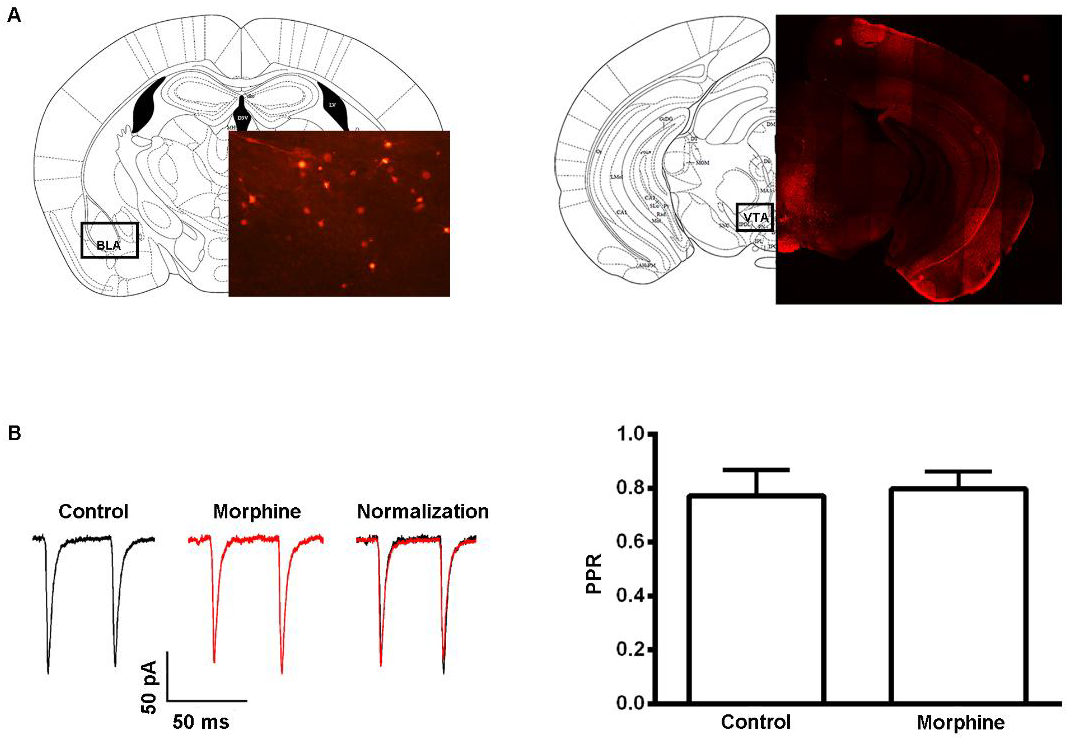
Effect of morphine on paired-pulse ratio (PPR) of light-evoked EPSC in DA neurons of the VTA from the BLA. (a) Left panel: image of coronal brain slice showing expression of ChR2-mCherry (red color) in the BLA at 6 weeks after the injection of the virus into the BLA. Right panel: Image of coronal brain slice showing expression of ChR2-mCherry in the VTA at 6 weeks after the injection of the virus into the BLA. (b) Left panel: typical original traces of the first and second EPSCs evoked by a pair of blue light pulses (470 nm, intervals of 50 ms at 0.1 Hz) before and after morphine (10 μM) as well as superimposed traces of the first and second EPSCs normalized to the amplitude of the first EPSC in DA neurons of the VTA. Right panel: average PPR evoked by a pair of blue light pulses (470 nm, intervals of 50 ms at 0.1 Hz) before and after morphine (10 μM) in VTA DA neurons from the BLA (n = 11 cells from eight mice, P > 0.05, compared to control before morphine).

**Figure 3.**
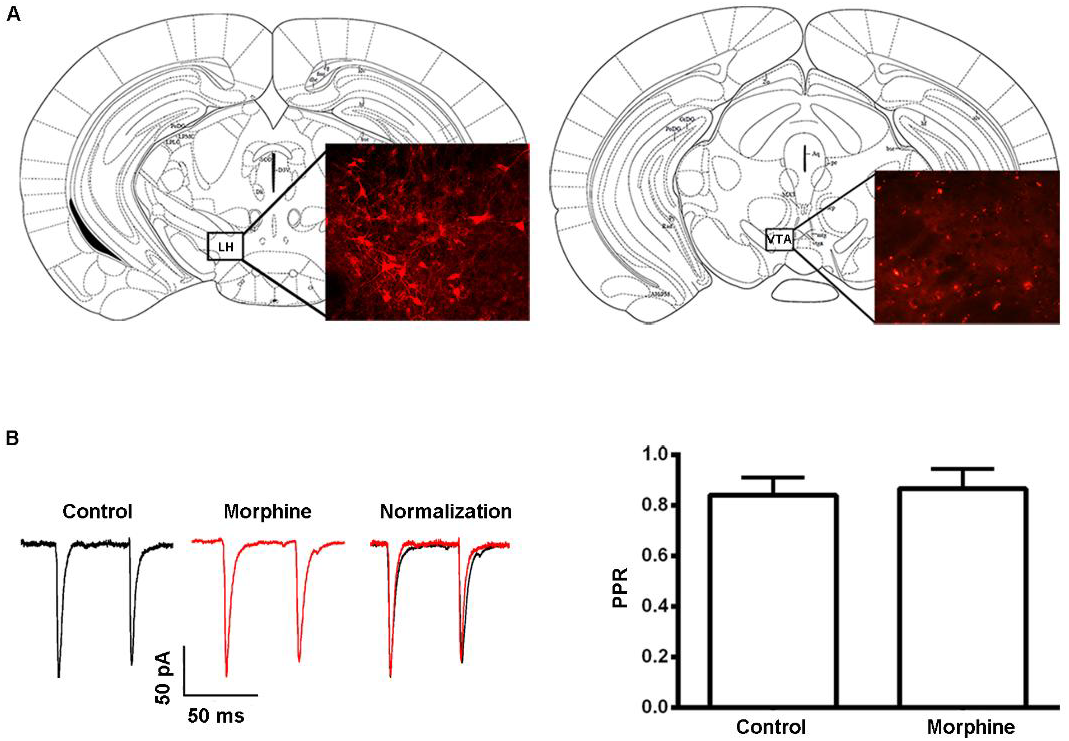
Effect of morphine on paired-pulse ratio (PPR) of light-evoked EPSC in DA neurons of the VTA from the LH. (a) Left panel: image of coronal brain slice showing expression of ChR2-mCherry (red color) in the LH at 6 weeks after the injection of the virus into the LH. Right panel: image of coronal brain slice showing expression of ChR2-mCherry in the VTA at 6 weeks after the injection of the virus into the LH (b) Left panel: typical original traces of the first and second EPSCs evoked by a pair of blue light pulses (470 nm, intervals of 50 ms at 0.1 Hz) before and after morphine (10 μM) as well as superimposed traces of the first and second EPSCs normalized to the amplitude of the first EPSC in DA neurons of the VTA. Right panel: average PPR evoked by a pair of blue light pulses (470 nm, intervals of 50 ms at 0.1 Hz) before and after morphine (10 μM) in VTA DA neurons from the LH (n = 7 cells from five mice, P > 0.05, compared to control before morphine).

### Inhibition of glutamatergic projection neurons from the mPFC to the VTA significantly reduces morphine-induced increase in locomotor activity of mice

To evaluate the contribution of morphine-induced increase in glutamate release from mPFC-VTA projection neurons to morphine-induced increase in locomotor activity of mice, we observed the influence of the inhibition of glutamatergic projection neurons from the mPFC to the VTA using chemogenetic method (Armbruster et al., 2007) on morphine-induced increase in locomotor activity of mice. We bilaterally injected a retrograde canine adenovirus expressing the Cre recombinase (CAV-GFP-Cre) into the VTA (Penzo et al., 2015), followed by the injection of the AAV-DIO-hM4Di-mCherry into the mPFC. Six weeks after the infection, the expression of AAV-DIO-hM4D(Gi)-mCherry was observed in the mPFC (left panel of Fig. 4a) due to retrograde transported CAV-GFP-Cre from the VTA (right panel of Fig. 4a) to the mPFC. This intersectional strategy can effectively target glutamatergic projection neurons from the mPFC to the VTA, which, subsequently, can be suppressed by treating mice with clozapine-N-oxide (CNO), the agonist of hM4Di. On this basis, we measured morphine-induced increase in locomotor activity of mice with and without CNO. The result showed that in control group where mice with hM4D(Gi) were given saline (i.p), morphine (10 mg/kg) could increase locomotor activity of mice. The average distance traveled in morphine+hM4D(Gi)+saline group was 305.4 ± 66.95 cm before and 4913 ± 392.6 cm during 5 min at 15 min after i.p morphine (10 mg/kg,n = 12 mice, two-way ANOVA, *F* _*(1,* *21)*_ *= 11.63*, ^*^P < 0.001, compared with control before morphine, left panel of Fig. 4b). However, in the CNO group where mice with hM4D(Gi) were given CNO (i.p), morphine-induced increase in locomotor activity of mice significantly decreased. The average distance traveled by mice in morphine+hM4D(Gi)+CNO group was 372.6 ± 51.67 cm before and 2973 ± 356.9 cm during 5 min at 15 min after i.p morphine (n = 11 mice, two-way ANOVA, *F* _*(1,* *21)*_ *= 11.63*, *P < 0.0001, compared with control before morphine; *F* _*(1,* *21)*_ *= 14.22*, ^#^P = 0.0011, compared with morphine+hM4D(Gi)+saline group, left panel of Fig. 4b). The average change percentage of distance traveled by mice in morphine+hM4D(Gi)+CNO was 964.1 ± 251.3 %, which was significantly lower than that (2551 ± 615.9 %) without CNO (independent t test, P = 0.0313, compared with morphine+hM4D(Gi)+saline group, right panel of Fig. 4b). This result suggests that morphine-induced increase in glutamate release from mPFC-VTA projection neurons contributes to morphine-induced increase in locomotor activity of mice.

**Figure 4.**
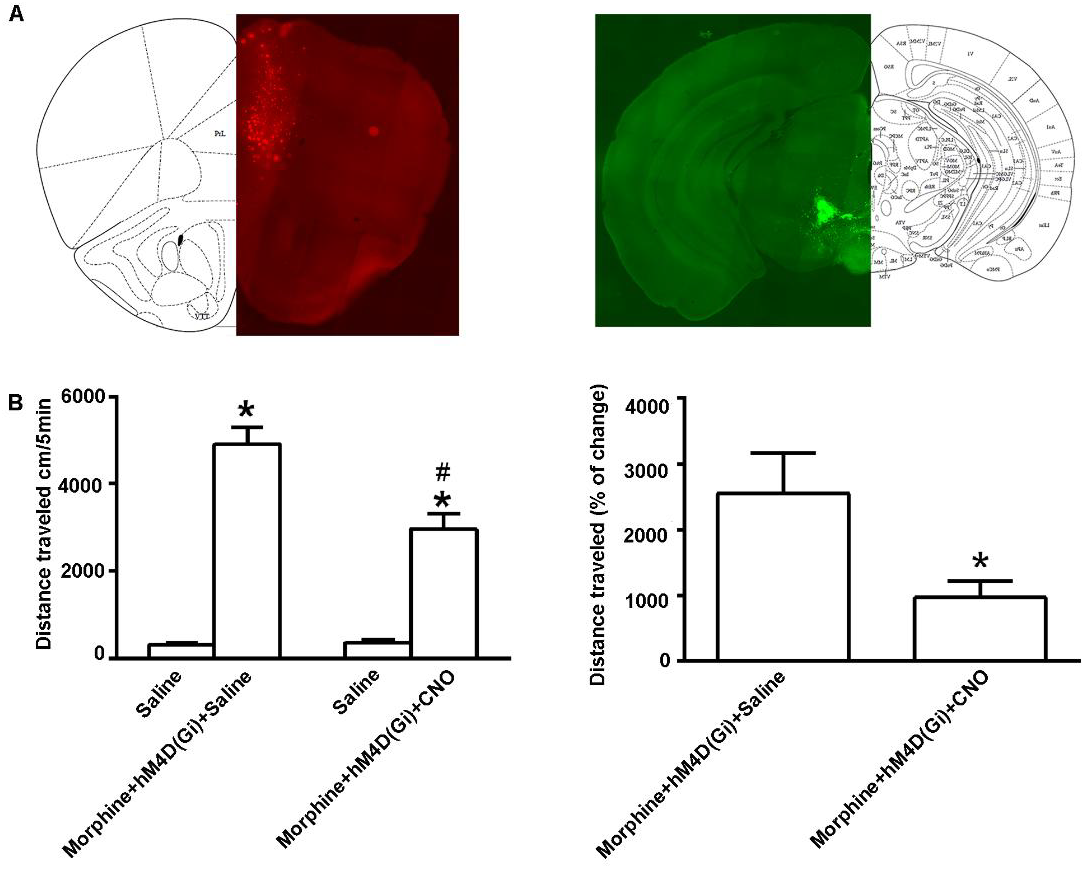
Influence of the inhibition of glutamatergic projection neurons from the mPFC to the VTA using chemogenetic method on morphine-induced increase in locomotor activity of mice. (a) Left panel: image of coronal brain slice showing AAV-hSyn-DIO-hM4D(Gi)-mCherry in the mPFC. Right panel: images of coronal brain slice showing CAV-GFP-cre in the VTA. (b) Left panel: average distance traveled by mice in morphine+hM4D(Gi)+saline (i.p, 0.2 ml/mice) group (n = 12, *P < 0.001, compared with control before morphine) and in morphine+hM4D(Gi)+CNO (i.p, 0.2 ml/mice group (n = 11, *P < 0.001, compared with control before morphine, ^#^P = 0.0016, compared with morphine+hM4D(Gi)+saline) during 5 min before and at 15 min after morphine (i.p, 10mg/kg). Right panel: average change percentage of distance traveled by mice in morphine+hM4D(Gi)+saline group (n = 12) and in morphine+hM4D(Gi)+CNO group (n = 11, P = 0.0313, compared with morphine+hM4D(Gi)+saline). Data are shown as the mean ± s.e.m.

## Discussion

It has been known that the activation of DA neurons of the VTA plays an important role in both natural stimulus and drug-induced reward response. Under natural state without drugs, direct optogenetic stimulation of VTA DA neurons has been demonstrated to be sufficient to modulate reward-related behaviors (Stuber et al., 2012). In further researches that involved how specific glutamatergic afferents modulate the activity of DA neurons in the VTA, it was found that the LH sent the largest subcortical glutamatergic projection to the VTA (Stuber et al., 2012) and electrical stimulation of the LH predominately increases the firing rates of VTA neurons (Stuber et al., 2012). This evidence suggests that LH-VTA glutamatergic projection neurons may be an important circuit in natural stimulus-induced reward. Another major source of glutamatergic input to the VTA comes from a long-range projection from the mPFC, which can target DA neurons of the VTA (Stuber et al., 2012). Stimulation of the mPFC led to the increase in extracellular glutamate in the VTA (Stuber et al., 2012), the activation of DA neurons of the VTA (Stuber et al., 2012) and the elevation of DA release in the forebrain (Stuber et al., 2012). These evidences suggest that glutamatergic projection neurons from the mPFC to VTA DA neurons may be another important circuit in natural stimulus-induced reward. In addition, DA neurons of the VTA also receive glutamatergic projections from the BLA (Stuber et al., 2012). However, for the drug-induced reward response, it is still unknown that morphine acts at what glutamatergic pathways to activate DA neurons in the VTA. Here, we find that morphine selectively promotes glutamate release from glutamatergic terminals of projection neurons from the mPFC to the VTA, but has no effect on that from the BLA or the LH to the VTA. This result suggests that morphine may mainly acts at glutamatergic terminals of projection neurons from the mPFC to the VTA to activate DA neurons of the VTA. This result is consistent with the report that the microinjection of morphine into the mPFC cannot produce rewarding effects (Liu et al., 2015) because the site of action of morphine on mPFC-VTA glutamatergic projection neurons is at terminals, rather than at the cell body in the mPFC.

What needs to be explained is that in our previous study in the VTA (Chen et al., 2015), when we use a pair of electrical pulses at intervals of 50 ms at 0.1 Hz to evoke EPSC, it always produces paired pulse facilitation, but here when we use a pair of optical pulses at intervals of 50 ms at 0.1 Hz to evoke EPSC in the VTA, it always produces paired pulse depression. This phenomenon also existed in the study by Vincent Pascoli et al (Pascoli et al., 2011). The reason for this difference remains unknown.

We further evaluate the contribution of morphine-induced increase in glutamate release of mPFC-VTA projection neurons to morphine-induced increase in locomotor activity of mice, which is an index of enhanced DA function in the VTA (Jalabert et al., 2011). The result shows that the inhibition of glutamatergic projection neurons from the mPFC to the VTA significantly reduces morphine-induced increase in locomotor activity of mice. This result was consistent with the report that the inactivation of the mPFC by the intra-mPFC injection of TTX could eliminate the acute morphine-induced excitation of DA neurons in the VTA (Liu et al., 2015).

## Materials and methods

### Virus injection

Bilateral injections of purified and concentrated pAAV-CaMKII-hChR2 (H134R)-mCherry, CAV-GFP-cre and AAV-hSyn-DIO-hM4D(Gi)-mCherry virus (2.05×10^12^ vector genomes/ml, Neuron Biotech Company, China) were stereotaxically performed in 6-week-old male C57/BL6 mice. Each side of the mPFC (final coordinates: AP, 1.95 mm; ML, ± 0.33 mm; DV, −2.4 mm from the skull surface), the BLA (final coordinates: AP, −1.4mm; ML, ± 3.0 mm; DV, −4.8 mm from skull surface) and the LH (final coordinates: AP, −2.8 mm; ML, ± 0.4 mm; DV, −4.2 mm from the skull surface) were respectively injected with 0.5 μl hChR2 (H134R)-mCherry. Each side of the mPFC (final coordinates: AP, 1.95 mm; ML, ± 0.33 mm; DV, −2.4 mm from the skull surface) was injected with 0.5 μl DIO-hM4D(Gi)-mCherry and each side of the VTA (final coordinates: AP, −3.08 mm; ML, ± 0.35 mm; DV, −4.8 mm from skull surface) was injected with 0.5 μl CAV-GFP-cre in the same manner (Chen et al., 2015). After the injection of virus, the animals were housed individually and were allowed to recover for over one week.

### VTA slice preparation

Male mice were anesthetized with chloral hydrate (400 mg/kg, i.p.). All experimental procedures conformed to Fudan University as well as international guidelines on the ethical use of animals. All efforts were made to minimize animal suffering and reduce the number of animals used. VTA slices were prepared according to procedures described previously (Hopf et al., 2007). The brain was removed rapidly from the skull and placed in modified ACSF containing 75 mM sucrose, 88 mM NaCl, 2.5 mM KCl, 1.25 mM NaH_2_PO_4_, 7 mM MgCl_2_, 0.5 mM CaCl_2_, 25 mM NaHCO_3_, and saturated with 95% O_2_ and 5% CO_2_ at ∼0°C. Horizontal 250 μm midbrain slices containing VTA were cut on a vibratome (VT-1200, Leica, Wetzlar, Germany) and transferred to normal ACSF containing 126 mM NaCl, 2.5 mM KCl, 1.25 mM NaH_2_PO_4_, 2 mM MgSO_4_, 2.5 mM CaCl_2_, 25 mM NaHCO_3_, and 10 mM glucose at 32°C. Slices were incubated for at least 60 min before patch-clamp recording.

### In vitro Optogenetics approach for electrophysiology

The medial terminal nucleus of the accessory optic tract (MT) was used as the anatomical structure to define the VTA (Hopf et al., 2007). VTA neurons were visualized on an upright microscope (BX50WI, Olympus, Tokyo, Japan) using infrared differential interference contrast or fluorescent optics. Whole cell current and voltage-clamp recordings were made using an EPC10 amplifier and PatchMaster 2.54 software (HEKA, Lambrecht, Germany). Electrodes had a resistance of 3–4 MΩ when filled with the patch pipette solution. The internal pipette solution contained 130 mM K-gluconate, 8 mM NaCl, 0.1 mM CaCl_2_, 0.6 mM EGTA, 2 mM Mg-ATP, 0.1 mM Na_3_-GTP, and 10 mM HEPES (pH 7.4). Cells were held at −70 mV under a voltage-clamp mode to record evoked EPSC. To observe PPR, two synaptic responses were evoked by flashing 470 nm light (5 ms pulses, 0.1 Hz), which delivered via an optical fiber (core diameter 200 μm, N.A. = 0.39, ThorLabs) coupled to a LED light source (Mightex) 500 μm above the recording cell. The series resistance (Rs) was monitored by measuring the instantaneous current in response to a 5 mV voltage step command. Rs compensation was not used, but cells where Rs changed by > 15% were discarded.

### Identification of VTA-DA neurons

After forming whole-cell recording mode, we identified DA neurons based on electrophysiological characteristics, which included both a spontaneous pacemaker-like firing and expression of a hyperpolarization-induced current (Ih) in the voltage-clamp configuration, by 1 s hyperpolarizing voltage steps (−70 mV to −150 mV) (Grace and Onn, 1989, Margolis et al., 2006, Zhang et al., 2010, Chieng et al., 2011).

### Locomotor behavior

The locomotor activity test was conducted as described previously with some modifications (Borgland et al., 2006). The locomotor activity of animals was monitored with a near infrared video camera within the operant chambers (Med Associates, St. Albans, USA). Distance traveled was measured using Open Field Activity Software (Med Associates) and analyzed locomotion estimates based on movement over a given distance and resting delays (movement in a given period of time). All animals were habituated to the test room for 2 h prior to the start of the experiment. Mice were habituated to the operant chambers for 5 min after 15 min intraperitoneal injection. After 15 minutes drug administration, the mice were placed in the chambers for the 5 min testing session. On days 1, 2, and 3, all mice were only given saline (intraperitoneal injections) to habituate them to the test protocol. On day 4, mice were given drugs according to experimental group. After the behavioral tests, all mice were anesthetized with an overdose of chloral hydrate and perfused with 0.9% saline. The brain was removed and fixed in 4% paraformaldehyde for 24 hr. Coronal sections (80 μm) were cut by a vibratome and fluorescence was observed under a microscope. Animals where the injection site was outside the VTA were discarded.

### Drug

K-gluconate, ATP·K_2_, GTP·Na_3_, 4-(2-hydroxyethyl) piperazine-1-ethanesulfonic acid (HEPES), ethyleneglycol-bis(b-aminoethyl ether) N,N,N’,N’-tetraacetic acid (EGTA), Triton X-100 and 0.01 M PBS were purchased from Sigma. Morphine was from Shenyang No.1 Pharmaceutical Factory, China. Clozapine-N-Oxide (CNO) was from Yihui Biological Technology Co, Shanghai, China. CNO was dissolved in 0.9% saline and administered at 10 mg/kg 30 min before the test (Ray et al., 2011, Li et al., 2013, Ray et al., 2013, Mahler et al., 2014).

## Data analysis

Numerical data were expressed as mean ± SE. Off-line data analysis was performed using a Mini Analysis Program (Synaptosoft), Clampfit (Axon Instruments), SigmaPlot (Jandel Scientific) and Origin (Microcal Software). Statistical significance was determined using Student’s t-test for comparisons between two groups or ANOVA followed by Student-Newman-Keuls test for comparisons among three or more groups, n refers to the number of cells. Every cell was from different slice and a group of cells in each experiment was from at least 5 animals. All post hoc comparisons were made using Tukey’s test. Results with P < 0.05 were accepted as being statistically significant.

## Acknowledgements

This work was supported by the National Program of Basic Research sponsored by the Ministry of Science and Technology of China (2009CB52201 and 2013CB835100), Science and Technology Program of Yunnan Province (2013GA003) and Project of Foundation of National Natural Science of China (31121061, 91332204, 81371466 and 31070932).p

